# XIBBIT: A biometric recognition tool for efficient *Xenopus laevis* identification and colony management

**DOI:** 10.64898/2026.06.30.735627

**Authors:** Dario Tomanin, Serban Tonie, Kerstin Bunte, Julia Kamenz

**Affiliations:** Molecular Systems Biology, Groningen Biomolecular Sciences and Biotechnology Institute, University of Groningen, 9747AG Groningen, The Netherlands; Bernoulli Institute for Mathematics, Computer Science and Artificial Intelligence, University of Groningen, 9700AK Groningen, The Netherlands

## Abstract

The African clawed frog *Xenopus laevis* is a widely utilized model organism in biomedical research; however, significant challenges in experimental reproducibility and colony management remain. A major obstacle lies in the reliable identification of individual animals, since frogs are generally housed in large groups and are difficult to distinguish due to their high morphological similarity. Conventional methods, including toe clipping and microchipping, are invasive and cause distress, emphasizing the need for non-invasive methods for accurate documentation and welfare monitoring.

In this study, we introduce XIBBIT (Xenopus Image-Based Biometric-pattern Identification Tool), a web-based application integrating computer vision and machine learning to identify individual *Xenopus laevis* based on their dorsal patterning. By exploiting these natural biometric signatures, the platform achieves reliable identification with up to 95.7% accuracy within three image captures under real life conditions.

In addition to identification, XIBBIT provides a centralized colony management system. It archives individual data, including health records and experimental histories, with customizable fields. To demonstrate XIBBIT’s capabilities, we used the application to track egg quality across repeated egg-laying events, revealing that egg quality is a repeatable, individual-specific trait in *Xenopus laevis*. Furthermore, we find seasonal effects on egg laying performance with the lowest performance during late-spring and summer months. Ultimately, XIBBIT provides an effective, time-efficient, and non-invasive solution to the problem of individual *Xenopus laevis* identification, facilitating both experimental reproducibility and high animal welfare standards.

## Introduction

The African clawed frog *Xenopus laevis* has long been a cornerstone of biomedical research, with hundreds of laboratories worldwide currently maintaining colonies of this species (1). *X. laevis* is valued for its broad utility across multiple disciplines, including cell and developmental biology, and as a model for a plethora of diseases, including cancer, congenital heart disease, spinal cord injury, and developmental diseases (2–5). Its practical and biological features, such as a long lifespan – some individuals can live 20 years or longer (6), straightforward husbandry, and key genetic similarities with humans (7), make it particularly well suited for studying fundamental biological processes and translating discoveries into insights relevant to human health. As with all animal models, effective colony management is essential for both animal welfare and scientific rigor. Group housing is often employed in *Xenopus* facilities to promote social interactions and reduce stress (8). However, this husbandry strategy introduces significant challenges for individualized tracking and record-keeping.

In research applications, oocytes surgically removed from the female frogs and egg extracts derived from *X. laevis* eggs are frequently used as experimental materials. The eggs are typically obtained by hormone-induced egg laying, a well-established technique that can be performed repeatedly in the same animal (3). However, egg quality can be influenced by multiple factors including water temperature, stress level, seasonality, genetic background, and reproductive history (9). Variations in these parameters directly affect experimental reliability and reproducibility (10). Robust individual identification of animals is therefore critical for correlating egg quality and reproductive performance with husbandry conditions, refining animal care, and improving scientific outcomes. Yet, the absence of non-invasive methods to track individual frogs has hindered systematic evaluation of these factors.

In addition to its importance for experimental reproducibility, reliable identification of *X. laevis* individuals has regulatory significance. Detailed records of the individual animals are essential for legal compliance with animal welfare legislation. By prioritizing the “Three Rs” principle of Refinement, researchers can reduce animal distress and improve welfare, ultimately increasing the scientific value and reproducibility of the study (11). Furthermore, *Xenopus laevis* has been designated an invasive species within the European Union, mandating stringent monitoring of the laboratory populations (12).

Conventional methods for identifying animals such as toe clipping, tattooing or microchipping remain problematic for *Xenopus* colonies (13, 14). These strategies are invasive, potentially painful and a source of discomfort for the animals. Moreover, frogs exhibit little external variation in appearance, which makes manual identification by caretakers and researchers difficult, especially when managing large colonies of several hundred animals. However, one distinguishing biological feature offers an opportunity: the dorsal skin of each frog displays a unique pattern of pigments (15). Like human fingerprints or cattle coat patterns (16) these markings serve as consistent and individualized biometric identifiers and provide a non-invasive and reliable basis for distinguishing individuals within a colony. Based on this notion, we previously developed a method that employs computer vision and machine learning to analyze dorsal markings of *X. laevis* frogs enabling automated and non-invasive recognition of individual animals (17). The approach provides a practical, welfare-oriented alternative to conventional methods, while simultaneously supporting reproducibility through the tracking of experimental and husbandry-related variables.

In the present study, we extend and formalize this approach with the development of XIBBIT (Xenopus Image-Based Biometric-pattern Identification Tool), a web-based application that integrates the dorsal-pattern recognition algorithm into a comprehensive colony management platform. XIBBIT can be simultaneously used as an identification tool and a centralized database, facilitating the storage and retrieval of relevant animal data, including health records, husbandry conditions, reproduction history, and experimental records. By consolidating information and enabling accurate and non-invasive identification, XIBBIT simplifies communication between researchers and caretakers, enhances reproducibility, and improves both efficiency and ethical standards of *Xenopus* colony management. Beyond these operational advantages, tracking the same animals across multiple egg-laying cycles allows researchers to investigate reproductive variables that were previously impossible to assess. As a proof of concept, we utilized XIBBIT to monitor reproductive performance over successive rounds of hormone-induced egg laying. This analysis suggests that egg quality does not fluctuate randomly between layings; rather, it constitutes a repeatable, individual-specific trait in *X. laevis*. Furthermore, the acquired longitudinal data suggests some degree of seasonal difference in the egg laying performance with the late spring and early summer months showing low performance.

## Results

### Design and implementation of the app

In order to make the developed identification algorithm (17) accessible to the broader scientific community, we developed XIBBIT, a web application that integrates machine learning-based dorsal pattern recognition into a user-friendly *Xenopus laevis* colony management platform. The backend handles all communication with the databases where animal and system information are stored; it also integrates the previously developed algorithm to train and apply pattern-based classification models for frog recognition. The frontend of XIBBIT was built using React (18) and displays a user-friendly application programming interface which provides the user with three main functions (Fig 1A):

1. Individual frog identification by submitting a photograph and retrieval of the corresponding experimental and welfare history. Users can subsequently modify, update, or delete frog records, e.g., change the location of a frog to a different tank or add information about a performed experimental procedure and experimental outcome.
2. Inquiry of general colony information, such as an overview of all frogs in the colony or the number and identity of the frogs inhabiting a specific tank.
3. General information of system’s parameters, e.g., daily monitoring of pH, temperature, and conductivity of the system’s water.

**Fig 1.**
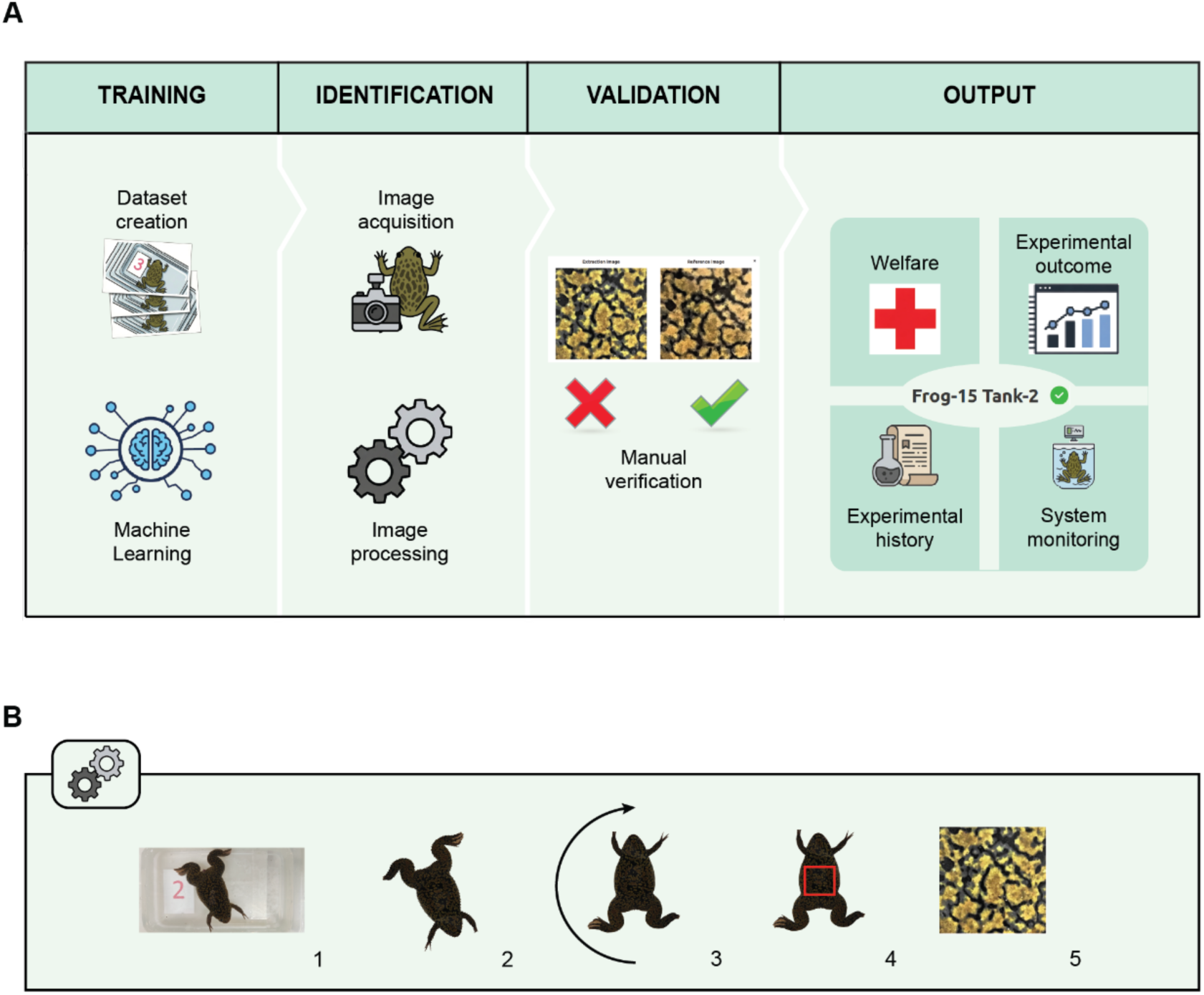
Schematic representation of XIBBIT. **(A)** Workflow for model training, animal identification, and databases usage. **(B)** Detailed workflow for the image processing performed by the algorithm. The shape of the frog is isolated from the background (1), the frog is rotated along its anterior-posterior axis into a vertical position (2), a region of the back is isolated (3) and its pattern extracted (4). The extracted pattern constitutes the basis for biometric pattern recognition.

To ensure data security, user authentication with privilege-based access is implemented. Users with basic privileges can view frog information, while those with higher-level privileges set by the administrator may add, edit, or delete frog records. User access is further restricted to the information of their respective institution or facility within the app. When setting up a new institution, the administrator has the opportunity to customize the web environment according to the specific operational requirements of the facility (e.g., number of tanks, system parameters to be monitored).

Recognition models are trained specifically for each facility. The system allows for the training of multiple models; if multiple models are available the user can manually select a model before uploading the photograph of the frog to be recognized. Once trained, the system identifies individual frogs from submitted images, either uploaded from a user’s device or captured directly through the app by granting access to the device’s camera. During processing, the algorithm isolates the frog from the background, rotates the image so that the head faces upward, and extracts the dorsal pattern (Fig 1B). Based on the extracted pattern, the system returns a predicted identity. Users verify the prediction by comparing the uploaded image with the reference image of the identified frog (Figs 1A and 2A). If the patterns match, the user can confirm and proceed to view or edit the animal’s information; if not, the identification can be rejected and a new image can be acquired and submitted. Each frog’s detail page provides customizable, editable fields for animal-specific data, such as the pictures of the animal (that the operator can save and use to monitor the animal’s condition over time), last injection date, welfare notes, egg quality, weight, and current tank number (Fig 2B). Action buttons include the standard *Save* and *Delete* options, an *Inject* button to update the hormone injection date, and a *Sick* button, which places a yellow exclamation mark beside the frog’s ID as a visual cue for closer monitoring during daily checkups (Fig 2C).

**Fig 2.**
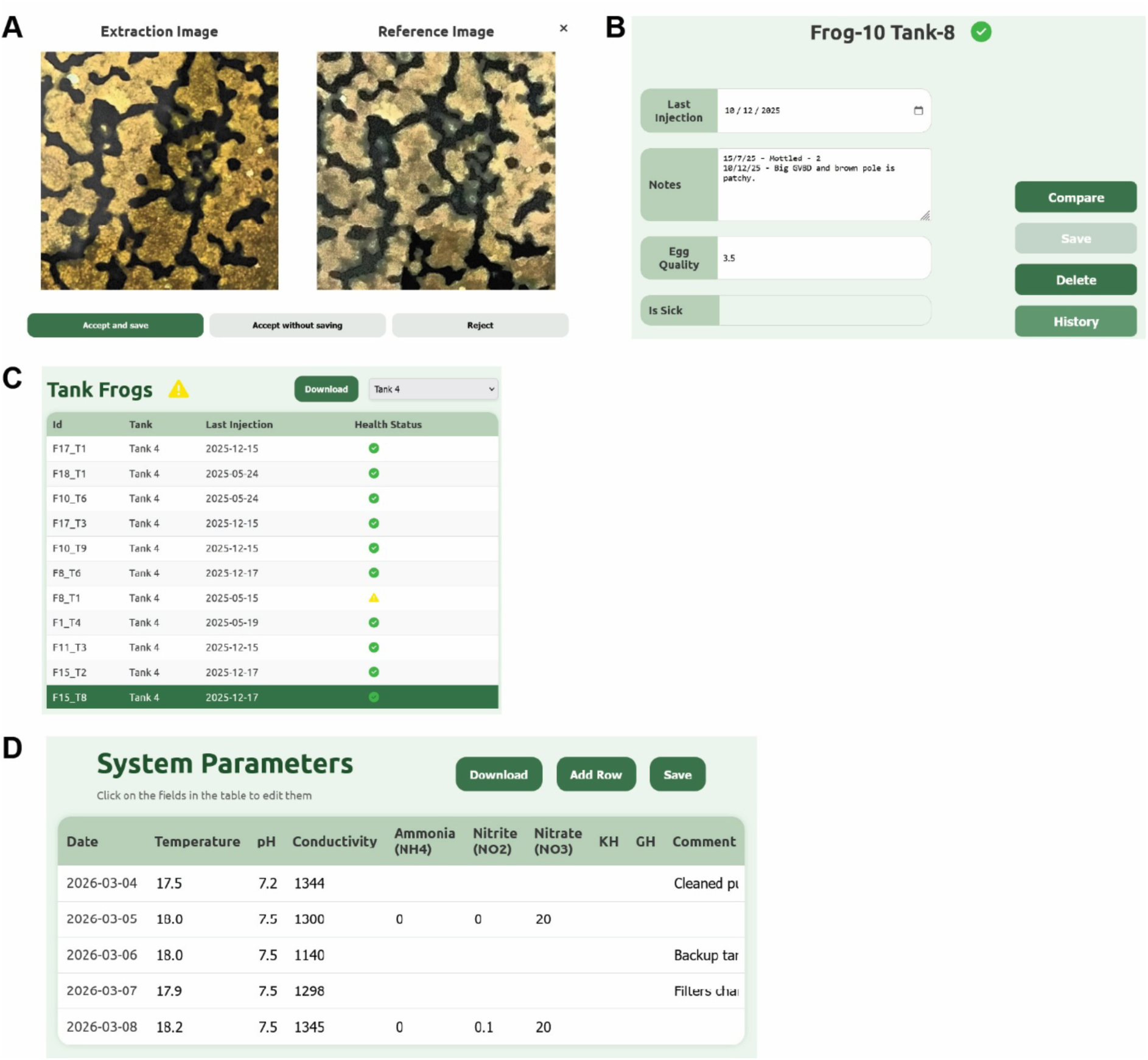
XIBBIT application interface. **(A)** Recognition interface showing a side-by-side comparison between the analyzed dorsal pattern and the reference pattern from the database. **(B)** Individual frog record page, showing editable fields for animal information, health status, egg quality, weight, and tank assignment. **(C)** Overview of the tank management page, detailing frog IDs, location, the date of the last injection and health status. **(D)** System parameters interface for recording system parameters of the aquatic system such as water temperature, conductivity, and pH.

The application maintains databases for animal welfare and experimental history, tank allocation, and system parameters, ensuring clear and organized data management for colony oversight and daily operations (Fig 2C). If necessary, this information can be exported as comma-separated value (CSV) files. The setup allows for comprehensive monitoring and management of all animals within a colony as well as system parameters critical to ensuring animal health (Fig 2D), such as water temperature and conductivity.

### Evaluation of frog identification efficiency

Previously, the recognition algorithm had been benchmarked by splitting the original dataset, consisting of 1647 images for 160 frogs, into training and test subsets, with all images acquired within the same 24-hour period (17). To create that dataset, we isolated the frogs in individual tanks and using a tripod acquired pictures in the morning (t_0h_), six hours later (t_6h_), and in the morning of the day after (t_24h_) using five different mobile devices. On that occasion, the model reached an accuracy up to 97.60%. However, for practical application of the algorithm and the associated app, it is critical to evaluate performance under more realistic conditions; for example, when the images of the animals to identify are collected months or even years after the acquisition of the original training set. To address this, we collected an additional dataset three years after the initial data collection, comprising 140 frogs that were also present in the original dataset. As in the original dataset (17), ten photographs were taken per frog using a tripod to maintain a fixed distance between the camera and the animal. Pattern extraction was performed unsupervised. The algorithm’s performance was assessed using two metrics: (1) the number of attempts required to correctly identify an individual, and (2) the overall percentage of correctly identified images within the dataset. To establish ground truth for the new dataset, we employed a semi-automated verification process with the final ID assigned only after human verification. This ensured that the evaluation metrics were based on 100% verified labels. Using tripod-supported photographs, 101 out of 140 frogs (72.1%) were correctly recognized on the first attempt. Of the remaining frogs, 18 (12.9%) were recognized on the second attempt, 12 (8.6%) on the third, and six frogs (4.3%) within four to ten attempts. Only three frogs out of 140 (2.1%) could not be identified within ten attempts (Fig 3A). Overall, 1003 out of 1399 images (71.7%) were correctly assigned to the corresponding individual (Fig 3B). Similar results were obtained by analyzing an additional dataset (Fig S1A).

**Fig 3.**
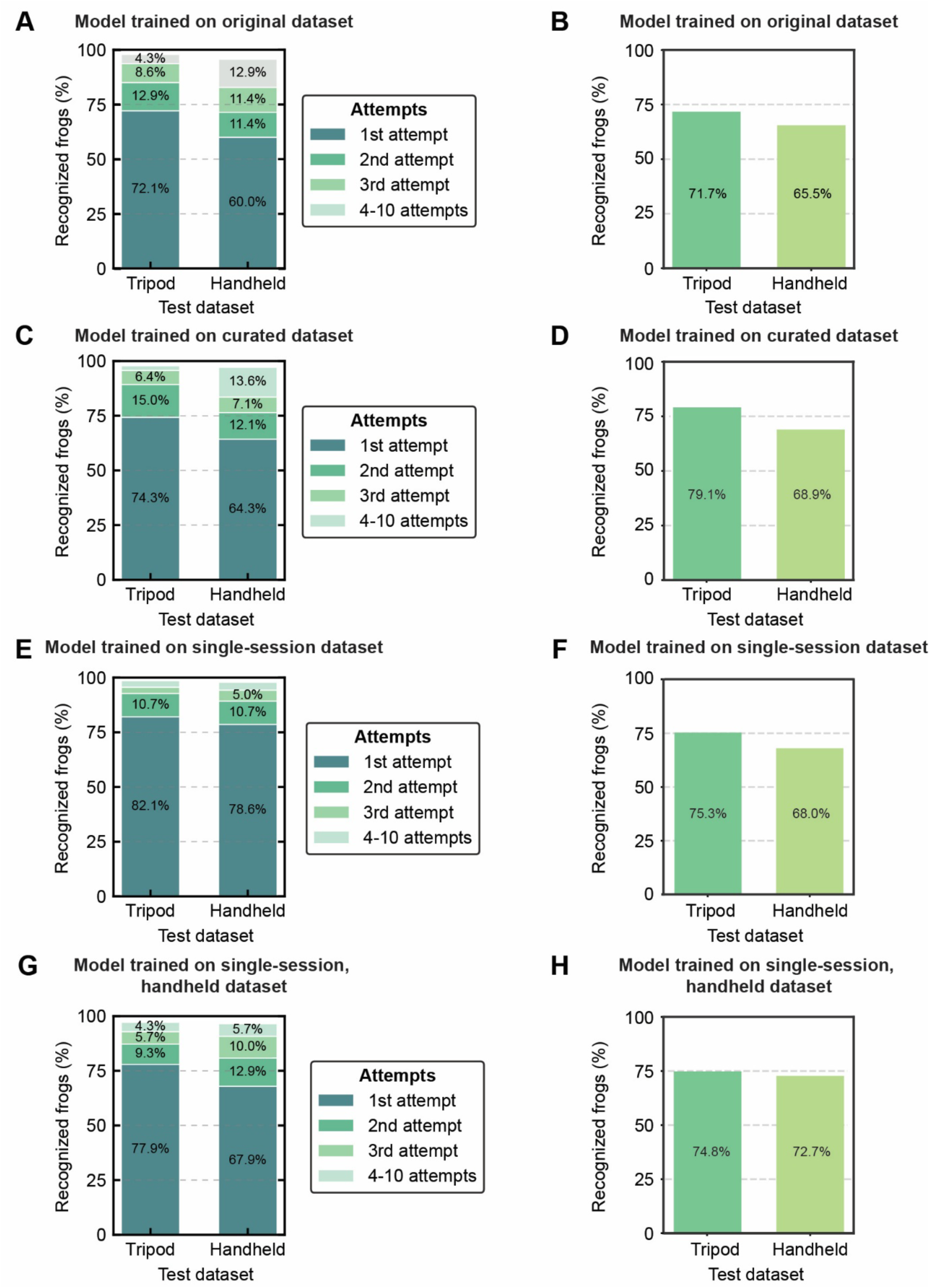
Accuracy of frog identification under different training and test scenarios. **(A)** Cumulative bar chart showing the percentage of frogs identified in one (dark green), within two, within three, or within ten attempts (progressively lighter shades of green) using the Original-trained model against the indicated single-session test datasets. **(B)** Overall recognition efficiency for model and test datasets in A. **(C)** Cumulative bar chart for the number of attempts required to correctly identify a frog using the model trained on the curated original dataset against the indicated single-session test datasets. **(D)** Overall recognition efficiency for model and test datasets in C. **(E)** Cumulative bar chart for the number of attempts required to correctly identify a frog using the Single session-trained model against the indicated single-session test datasets. **(F)** Overall recognition efficiency for model and test datasets in E. **(G)** Cumulative bar chart for the number of attempts required to correctly identify a frog using the Handheld-trained model against the indicated single-session test datasets. **(H)** Overall recognition efficiency for model and test datasets in G.

As tripod-assisted image acquisition is laborious and time-consuming, we also assessed the algorithm’s performance, when images were acquired simply using a handheld device without any static support (Handheld dataset). Again, ten images per individual were acquired and blurry images were removed from the set. Under these less standardized conditions, 84 frogs (60.0%) were recognized on the first attempt, 16 frogs (11.4%) on the second attempt, and 16 frogs (11.4%) on the third attempt. Of the remaining animals, 18 frogs (12.9%) were recognized within ten tries, whereas 6 frogs (4.3%) were not identified (Fig 3A). Overall, 903 out of 1379 images (65.5%) were detected using this model (Fig 3B). The lower performance can in part be explained by a higher failure of extracting the correct region of the dorsal pattern, along with variability in lighting conditions, camera shake, and subtle motion blur that degraded pattern features despite manual removal of overtly blurry photos. These factors can alter local contrast, sharpness, and texture statistics, thereby challenging the model’s feature matching and highlighting opportunities for improvements in future releases.

### Impact of dataset acquisition protocols on identification accuracy

To improve identification accuracy, we first examined the 1647 images used to train the original model and found that the dataset included numerous images that were either blurry, out of focus, improperly rotated (>90°), or where the extraction algorithm extracted the wrong area of the frog’s back, sometimes even including background elements. We removed 148 such images obtaining a new training dataset consisting of 1499 pictures. We then trained a new model using this dataset, which we will refer to as the ‘curated’ dataset, to assess the effect of excluding flawed images.

As a result of this curation, recognition rates improved modestly for test images acquired with tripod support: 104 frogs (74.3%) were recognized on the first attempt, 21 (15.0%) on the second, nine (6.4%) on the third, four (2.9%) within four to ten attempts, and only two (1.4%) were not recognized during this experiment (Fig 3C). Overall, 1106 images out of 1399 (79.1%) were correctly assigned (Fig 3D). This represents a 7.4% improvement in accuracy compared to the model trained with the original non-curated dataset, underscoring the critical role of high-quality training data. Similarly, the model’s ability to recognize handheld-acquired test images improved: 90 frogs (64.3%) were recognized on the first attempt, 17 (12.1%) on the second, 10 (7.1%) on the third, 19 (13.6%) within four to ten attempts, and only four (2.9%) were not recognized (Fig 3C). Overall, 950 handheld-acquired pictures out of 1379 were identified (68.9%) compared to 903 out of 1379 images (65.5%) (Fig 3D).

The original training data had been acquired by taking ten photographs over a period of 24 hours using different devices and a tripod, accounting for changes in skin pigmentation as a result of different environments, differences in camera quality, and ensuring a defined distance to the animal. However, this approach is time-consuming and logistically challenging for large colonies. Hence, we evaluated whether a training dataset collected at a single time point could suffice, eliminating the need to use multiple devices and staggered time points. Acquiring pictures in a single session also avoids the necessity to house frogs individually for the period of data acquisition, thus reducing animal discomfort. Using a model trained on such single-session dataset acquired using a tripod performed competitively compared to the curated multi-session dataset: analyzing the tripod-acquired test set, 115 frogs (82.1%) were recognized on the first attempt, 15 (10.7%) on the second, four (2.9%) on the third, four (2.9%) within four to ten attempts, and only two (1.4%) were not recognized at all (Fig 3E). In total, 1011 out of 1343 images (75.3%) were recognized (Fig 3F). When testing the handheld test set with this model, 938 out of 1379 images (68.0%) were paired with their correct ID (Fig 3F). 110 frogs (78.6%) were recognized on the first attempt, 15 (10.7%) on the second, seven (5.0%) on the third, five (3.6%) within four to ten attempts, and three (2.1%) were not recognized (Fig 3E). While some improvements might be due to the training and test datasets being acquired within a few months of each other, the results nevertheless demonstrate that a model trained on data from a single time point performs comparably to one trained with color adaptation in mind.

Since using a tripod may not always be practical, we also trained a model using a single-session dataset acquired entirely without support. The model trained with this dataset demonstrated strong performance, closely approaching the results of the tripod-assisted single-session trained benchmark: for the single-session test set with tripod support, 109 frogs (77.9%) were recognized on the first attempt, compared to 95 frogs (67.9%) for the handheld test set (Fig 3G). Similarly, while 138 frogs (98.6%) were eventually recognized in the single-session set, the handheld set followed closely with 135 frogs (96.4%) identified (Fig 3H). Using this handheld-trained model, 74.3% of the single-session tripod-mounted dataset pictures have been correctly labelled (998/1343) (Fig 3H), while 1003 out of 1379 handheld images (72.7%) have been recognized (Fig 3H).

### Analysis of the robustness of the training conditions

To evaluate the robustness of models trained on a data set acquired in a single session against variations in camera hardware and environmental conditions (such as skin color adaptation), we tested them using the original, curated dataset as a benchmark, which was acquired with several different devices and over a time span of 24 hours.

While the model trained on the curated original dataset previously achieved a 79.1% recognition rate (1106/1399) against a test data set acquired in a single session, the opposite was not the case. The model trained on data acquired within a single session significantly underperformed when the conditions were reversed. Specifically, the model achieved a recognition rate of only 41.8% (550/1317) when analyzing the more diverse images of the curated original dataset (Fig 4A). This significant decrease highlights the challenge of identifying frogs when the training data lacks the environmental and physiological variability present in the test set (Fig 4A).

**Fig 4.**
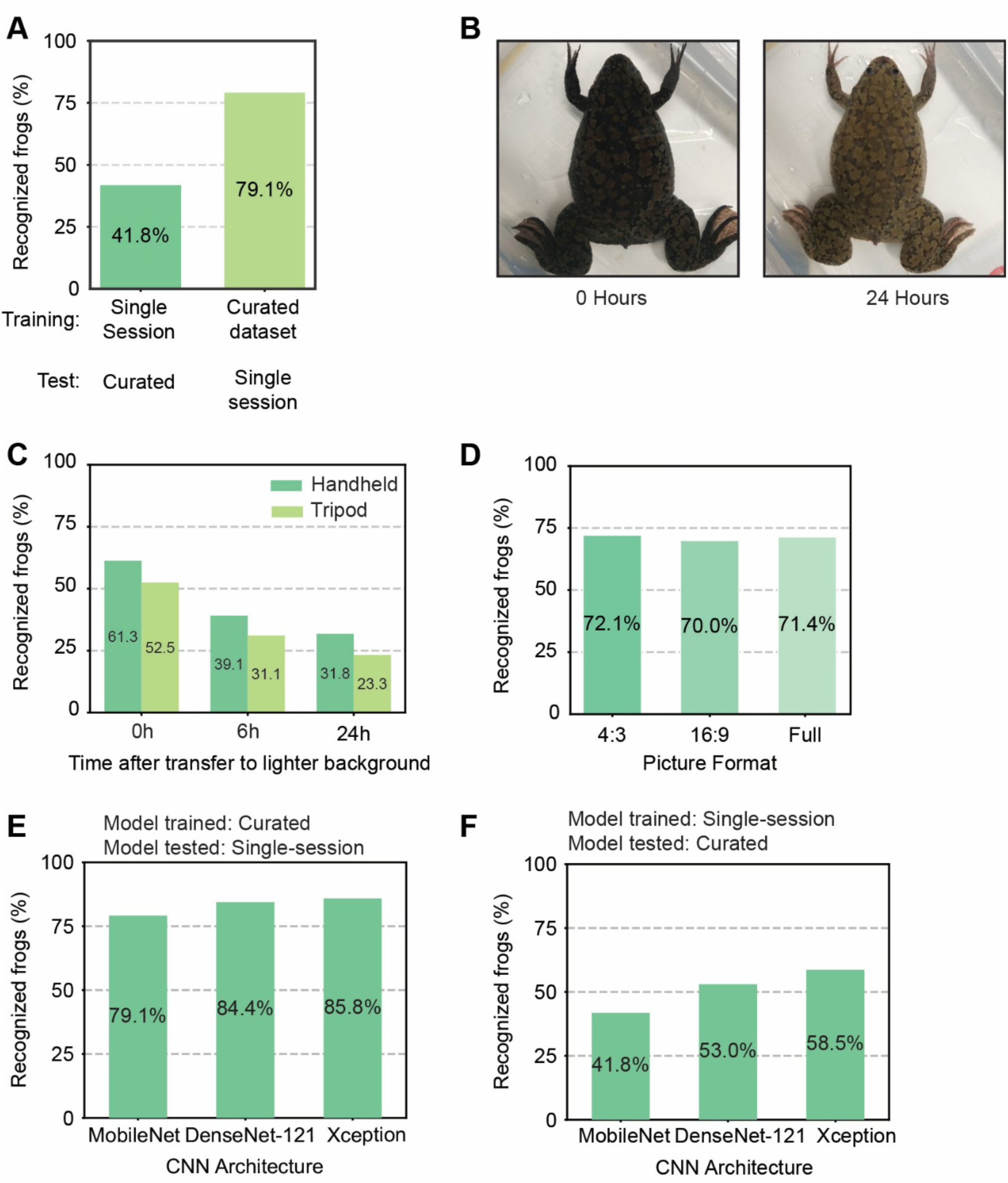
Robustness of frog identification against color adaptation, image format, and model architecture. **(A)** Overall image recognition rate of a model trained on the single-session dataset tested against the curated dataset, compared to a model trained on the curated dataset tested against the single-session dataset. **(B)** Representative photograph of the same frog before (t_0_) and after color adaptation (t_24_). **(C)** Bar chart representing the overall recognition efficiency of models trained on a single-session handheld dataset (dark green) or a single-session tripod dataset (light green) tested against the curated dataset divided by acquisition time point at 0, 6, and 24 hours (t_0_, t_6_ and t_24_, respectively) after placing the frog in a container with a lighter background. **(D)** Bar chart showing the recognition efficiency of the curated-trained model across three image acquisition formats (4:3: 4096 × 3072 px; 16:9: 4096 × 2304 px; full screen: 4096 × 1856 px), acquired using a tripod-mounted smartphone. **(E)** Bar chart representing the overall recognition rates of MobileNet, DenseNet-121, and Xception architectures using a model trained on the curated dataset and tested on the single-session dataset. **(F)** Bar chart showing the overall recognition rates of MobileNet, DenseNet-121, and Xception architectures using a model trained on the single-session dataset and tested on the curated dataset.

This discrepancy in model efficiency can be explained by the different heterogeneity of the two training datasets. The original curated dataset included pictures of frogs acquired using different devices and at different time points (0, 6, and 24 hours) after removing the frogs from an environment with a dark background and placing them into a tank with a lighter background. During this 24-hour period, the contrast and color intensity of the frogs’ dorsal patterns changed significantly (Fig 4B). Conversely, the single session dataset only included pictures acquired at a single timepoint and with a single device resulting in a less heterogeneous dataset. Indeed, when the original curated dataset was stratified by acquisition time (0 h, 6 h, and 24 h after moving the frogs to a lighter background), about 30% more images were recognized at 0 h than at 24 h (Fig 4C). In addition, across all time points, accuracy was approximately 8% higher when using a model trained on a single-session handheld dataset than one trained with tripod-based support (Fig 4C).

We also addressed whether the image format used during picture acquisition affected frog recognition. Three photographs per frog were sequentially acquired using a tripod-mounted smart phone in 4:3 (4096 px × 3072 px), 16:9 (4096 px × 2304 px), and full screen (4096 px × 1856 px) format, respectively. Out of 143 frogs tested, 101 (72.1%) were recognized using the 4:3 format, 98 (70.0%) using 16:9, and 100 (71.4%) using full screen format (Fig 4D). In this test, approximately 70% of frogs were identified with a single picture, and around 85% were recognized within three attempts across the different formats. This result indicates that picture format has a negligible impact on identification accuracy, demonstrating that the recognition algorithm is robust to format conversions.

Taken together these data highlight that while the identification algorithm is resilient to image format and aspect ratio variations, its performance is critically dependent on the heterogeneity of the training data. The dramatic drop in accuracy when moving from controlled to diverse environments suggests that physiological color adaptation and variability in light conditions are the primary bottlenecks for cross-session reliability.

### Convolutional Neural Network efficiency

Originally, XIBBIT was designed to run locally on user smartphones using the lightweight MobileNet architecture (19). As we started hosting the application on a server, the lightness of the architecture became a secondary aspect; for this reason, we decided to compare the average recognition rates of the models based on MobileNet, with those with DenseNet-121 (20) and Xception (21) architectures. Using the curated original dataset, the Xception architecture achieved the highest accuracy at 85.8% (1200/1399 images). This performance was comparable to that of DenseNet-121, which reached 84.4% (1180/1399), followed closely by MobileNet with an accuracy of 79.1% (1106/1399) (Fig 4E).

A similar performance hierarchy was observed when the models were trained on the single-session dataset and tested against the curated dataset. In this scenario, Xception achieved the highest recognition rate at 58.5% (771/1317 images), followed by DenseNet-121 at 53.0% (697/1317 images), while MobileNet dropped to 41.8% (550/1317 images) (Fig 4F). Overall, Xception outperformed the other architectures across both conditions, while MobileNet showed the steepest decline when generalizing to data outside its training distribution. The substantial drop in accuracy observed for all models when trained on the single-session dataset and tested against the more heterogenous curated original dataset underscores the critical role of diversity in the trainings data for achieving robust, generalizable recognition performance. These results highlight opportunities for further improvements by choosing a less lightweight architecture for model training.

### Egg quality is a moderately stable individual trait

To evaluate the utility of XIBBIT for longitudinal tracking of individuals and reproductive outcomes, we used the platform to record egg quality across repeated hormone-induced egg-laying events in our colony. Thus far, egg quality has been assessed for 249 egg-laying events from 114 individual frogs, including 32 animals with records from three consecutive laying events. After each hormone induction, eggs from the identified frog were scored on a scale of 0–4 (0 = no laying or unusable eggs; 4 = excellent egg quality). Scores were assigned based on overall yield, the proportion of apoptotic eggs, the presence of immature oocytes (evidenced by stringiness or clumped eggs), egg pole pigmentation, and the distinctness of the white spot marking germinal vesicle breakdown.

Based on these scores, frogs were classified into three categories: good layers (n = 14), with a mean egg quality score >2.5 and low variability across egg-laying events (standard deviation <0.6); unreliable layers (n = 13), with mean scores <2.5 and generally higher variability in egg quality between events (>1 for 10 out of 13 frogs); and non-layers (n = 5), which consistently failed to produce eggs or produced eggs of extremely poor quality (Figs 5A and 5B). To quantify the consistency of egg laying performance, we calculated the intraclass correlation coefficient (ICC) for these 32 frogs. The calculated ICC was 0.63 (95% CI: 0.44-0.78; F(31, 64) = 6.02, p < 0.001), indicating that approximately 63% of the total variation in egg quality is attributable to stable between-individual differences rather than to variability across multiple layings of the same frog. Even at the lower bound of the confidence interval (ICC = 0.44), individual identity remains a meaningful source of variation, suggesting that egg-laying performance is a genuine, repeatable trait rather than a purely stochastic outcome.

**Fig 5.**
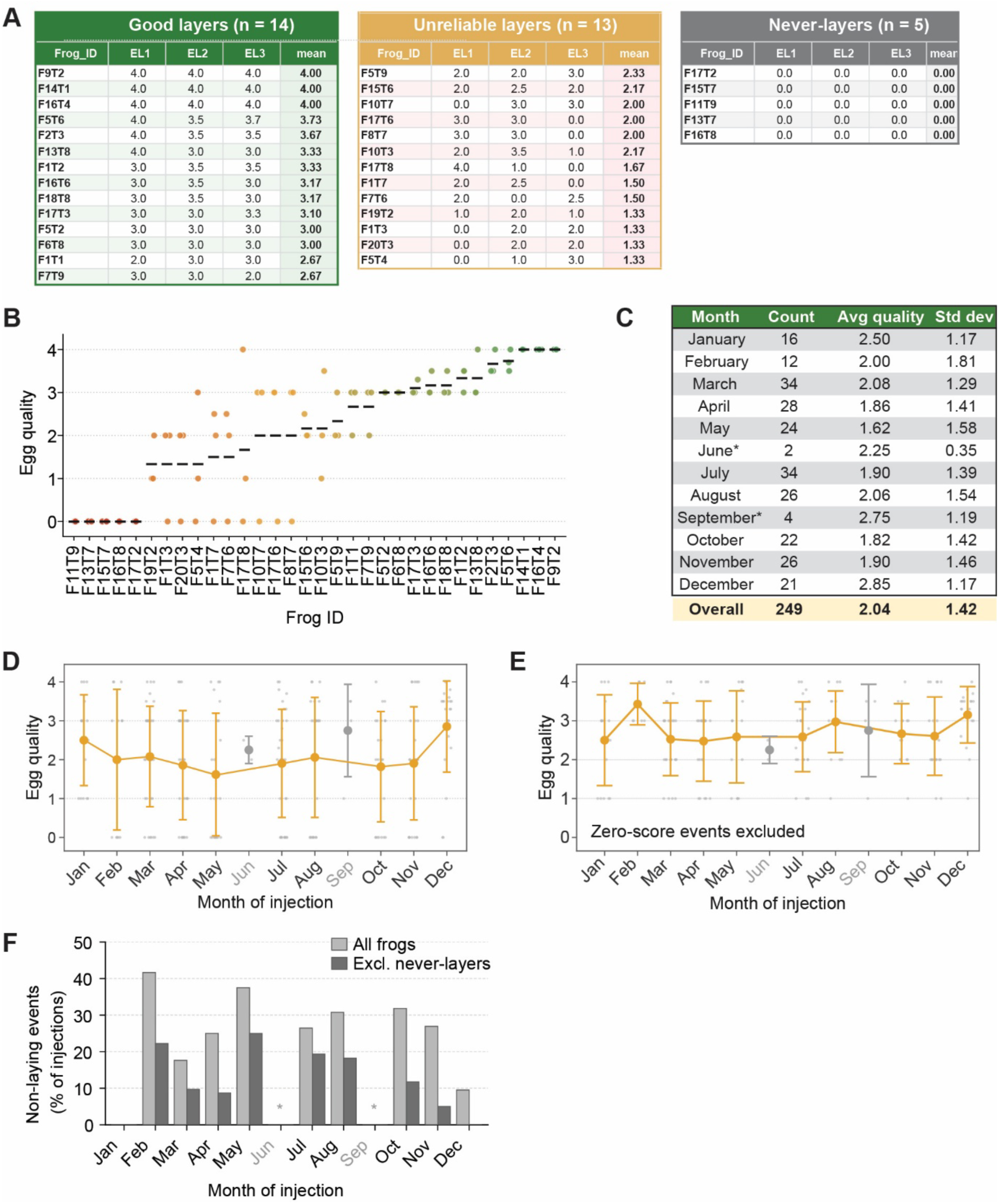
Egg quality is a repeatable individual trait in Xenopus laevis. **(A)** Egg quality scores of the 32 frogs for which three consecutive hormone-induced egg-laying events (EL1 - EL3) were recorded. On the basis of their mean score, individuals are grouped into good layers (mean ≥ 2.5; n = 14; green), unreliable layers (0 < mean < 2.5; n = 13; red), and non-layers, which never produced usable eggs (mean = 0; n = 5; grey); the mean of the three events is given for each frog. **(B)** Egg quality of 32 individual frogs, each assessed across three independent egg-laying events. Each point represents the quality of a single laying (scored 0–4; three layings per frog), and the black bar denotes the per-frog mean. Frogs are ordered along the x-axis by ascending mean egg quality; point and bar color encode this mean on a gradient from low (red) to high (green). **(C)** Average egg quality per month of injection, reporting for each month the number of recorded scores (count), the average of the quality, and the standard deviation, together with the overall values computed across all injections (n = 249). Months marked with an asterisk are based on few recorded injections (June, n = 2; September, n = 4) **(D)** Mean egg quality (± standard deviation) for each month of injection, corresponding to the data tabulated in (C). Months marked with an asterisk are based on few recorded injections (June, n = 2; September, n = 4). **(E)** Mean egg quality (± standard deviation) for each month of injection excluding non-laying events, corresponding to the data tabulated in (C). Months marked with an asterisk are based on few recorded injections (June, n = 2; September, n = 4). **(F)** Percentage of hormone injections that resulted in a non-laying event per month. Light grey bars include all frogs (60 zero scores across 249 injections); dark grey bars exclude the 16 frogs that never produced usable eggs in any recorded injection. Asterisks mark months based on very few injections (June, n = 2; September, n = 4).

Seasonal variations in oocyte quality have been reported previously (22), including seasonal variations in the quality of laid eggs in temperature-controlled environments (23). Stratifying our longitudinal data by the month in which the egg laying occurred indeed revealed seasonal differences (Figs 5C and 5D). The winter months, December and January, exhibited the highest mean scores (2.85 and 2.5, respectively), whereas May yielded the lowest (1.62). June and September had too few egg-laying events to be reliably evaluated. Omitting zero-score events, i.e. events where no eggs or extremely poor eggs were laid, largely abolished the seasonal differences (Fig 5E). Consistent with this notion, the rate of non-laying events varied across the year. Whereas non-laying events were rarely observed in the winter months (0 out of 16 in January, 2 out of 21 in December), more than 1 out of 4 injections did not yield eggs in May, July, and August. The trend was even more pronounced when the never-layers were excluded from this analysis (Fig 5F). February constituted an exception to this trend, likely because of the overall low number of total injections performed and large number of never-layers injected during that month.

Whereas the seasonal trends even under controlled environmental conditions requires further confirmation in larger datasets, the observed pattern is consistent with reports of seasonal variation in *Xenopus* oocyte and egg quality, which persists even in temperature-controlled laboratory colonies (22, 23). The underlying basis is not firmly established, however we could not find any other significant correlations between egg quality and monitored system parameters (e.g., temperature, pH) at least not within the variation that occur in the tightly controlled environment of our system (Fig S2).

Taken together, these results suggest that egg quality *in Xenopus laevis* is shaped by both stable inter-individual differences and a seasonal component. More importantly, they illustrate that being able to monitor these sources of variation is only possible when individual egg layings can be attributed to specific animals over extended periods. A reliable, non-invasive identification system such as XIBBIT is therefore an excellent tool for systematically monitoring and exploiting this variation, for example by preferentially selecting consistent good layers, scheduling injections during favourable months, and excluding persistent never-layers, thereby improving experimental planning, reducing animal use, and enhancing reproducibility.

## Availability and future directions

The development and implementation of XIBBIT represents a substantial advancement in the management of *Xenopus laevis* colonies within research environments. Our results demonstrate that this non-invasive, pattern-recognition system provides a reliable alternative to traditional identification methods, such as toe clipping or microchipping, which are known to negatively impact animal welfare. By eliminating the necessity for physical intervention, XIBBIT adheres to the refinement principle of the Three Rs framework, enhancing welfare standards without imposing excessive procedural demands on personnel.

The system demonstrated high practical utility, achieving an identification accuracy of approximately 70% on the first attempt and typically exceeding 85% within three attempts, with rates reaching over 90% under optimized training conditions (summarized in Figs S1B-E). To address accuracy challenges, the system includes immediate verification against the reference individual. The immediate feedback allows users to instantly recapture an additional picture if necessary. This iterative process ensures accurate identification with minimal time investment. Furthermore, the algorithm exhibited technical robustness across various image formats, including 4:3, 16:9, and full-screen aspect ratios, thereby accommodating hardware variations and reducing the need for rigid imaging protocols.

Optimization of training protocols significantly improved the system’s performance. Specifically, the removal of poor-quality images and the use of tripod-assisted data collection enhanced recognition accuracy while minimizing the duration for which animals were separated from their colonies. While handheld image acquisition resulted in slightly lower performance compared to stable supports, the success rates remained acceptable for most applications. These findings suggest that while the system is adaptable to less controlled settings, a strategic approach is optimal: utilizing stable supports for generating training datasets and handheld acquisition for routine, day-to-day identification. Only if changes in color intensity and contrast are anticipated, a training dataset accounting for such changes will be required.

The selection of model architecture had a modest impact on accuracy in controlled settings, where Xception and DenseNet-121 outperformed MobileNet by approximately 6% on the single-session dataset and by more than 10% for a more heterogenous dataset. This suggests that whereas MobileNet remains competitive under more homogeneous conditions, deeper architectures such as Xception and DenseNet-121 generalize better to datasets encompassing greater variability.

Beyond identification, the integrated database (linking individual IDs to health, husbandry, and experimental metadata) enables centralized record-keeping and improves communication between researchers and facility staff. Dedicated tools, such as the “Sick” and “Inject” cues, ensure that critical welfare and experimental data are immediately accessible through a secure interface. The application also allows the saving of additional images along the way, providing the possibility to retrain the models regularly with a larger training set. Despite these advancements, several opportunities for improvement remain. A small proportion of frogs could not be identified within ten attempts, indicating a need for continued improvement in image quality control and robustness of the machine learning models. Additionally, while XIBBIT shows significant potential for broader use, its performance in recognizing other Xenopus species, e.g. *Xenopus tropicalis,* and other applications (e.g., field studies) has not formally been evaluated. Existing tools for non-invasive amphibian identification fall into two groups, and XIBBIT differs from both. The classical tools (Wild-ID, I3S Pattern+, APHIS, AmphIdent and ManderMatcher) match images pairwise, either by analyzing specific features or by pixel cross-correlation. These tools then return a ranked list of candidates for the user to confirm. These kinds of tools typically require manual pre-processing such as cropping or keypoint placement (24, 25). The newer, deep-learning systems instead identify an animal by comparing its pattern to those in a database and finding the most similar one, and they can also flag individuals that are not yet registered (26). This open-set design is well suited to wild populations, where new individuals continually appear, whereas in a managed colony the set of animals is known and controlled, making closed-set classification the more appropriate choice. XIBBIT differs from both: it identifies frogs with a trained convolutional neural network that assigns each image directly to an enrolled individual (closed-set classification), through a fully automated pipeline that requires no manual pre-processing. Nevertheless, XIBBIT constitutes the first widely available, non-invasive and accurate methodology for the management of *Xenopus laevis* colonies. It offers distinct advantages regarding animal welfare, data reproducibility, and facility oversight, establishing a foundation for further technological integration in amphibian research.

XIBBIT is accessible as a web application hosted by our university (https://frog.web.rug.nl/). Alternatively, the source code is publicly available via https://zenodo.org/records/19552231.

## Methods

### Animal Handling and Preparation for Imaging

The University of Groningen is housing a colony of 140 female *Xenopus laevis* in accordance with national animal welfare laws and reviewed by the Animal Ethics Committee of the Royal Netherlands Academy of Arts and Sciences (KNAW) under a project license granted by the Central Committee Animal Experimentation (CCD) of the Dutch government and approved by the University of Groningen (IvD), with project license number AVD 10500202114408. The colony is divided into groups of ten to twenty individuals per aquatic tank.

### Image Acquisition

For image capture, each frog was placed in an open transparent water container on a white background. Depending on the experimental question, one of five different acquisition protocols was used. For the original dataset, each frog was photographed (ten images per frog) across three different timepoints (0 hour, 6 hours, and 24 hours) using different smartphone devices, after removing them from a dark background environment and placing them into a lighter background environment, as previously described (17).

The curated training dataset was obtained by manually removing images with suboptimal extraction based on predefined criteria, including blurring, dorsal pattern rotation greater than 90°, or inconsistent body positioning within the extraction window.

In contrast to the “original” dataset, where frogs were photographed over a 24-hour period, the single-session dataset was captured within a short time period. For this protocol, each frog was photographed ten times in rapid succession, capturing the animal in different positions using a smartphone mounted on a tripod above the tank. Similarly, for the “handheld” dataset, ten images per frog were captured in a single session with the smartphone held freehand. Finally, to evaluate the impact of image format, an additional dataset was acquired where each frog was photographed three times using the tripod setup at different aspect ratios: 4:3 (4096 × 3072 pixels), 16:9 (4096 × 2304 pixels), and full-screen mode (4096 × 1856 pixels).

### Identification Procedure

XIBBIT’s classification model was trained on the corresponding dataset for each experimental condition using the MobileNet architecture if not otherwise stated. Once the models were trained, the application was used to identify individual frogs across independent test sets, performing recognition analysis on the ten available test images for each subject. The identification process involved sequentially submitting each test image to the application and recording the number of attempts required for correct identification. Recognition was defined as successful if the correct identity was returned by the application within ten sequential image submissions. For each frog, the minimum number of images required for successful recognition was recorded. If a frog was not identified correctly within ten attempts, it was scored as a failed recognition. Identification results were manually verified by reviewing the comparison output provided by the application’s interface, ensuring correspondence with the true frog identity.

### Egg quality analysis and repeatability

After each hormone-induced egg-laying event, the eggs of the identified frog were inspected and evaluated on a scale from 0 to 4, where 0 indicates that no eggs were laid or that the eggs were unusable and 4 indicates excellent eggs. The score was assigned on the basis of three different criteria: the proportion of apoptotic eggs, the presence of immature oocytes (“stringiness”), and the color of the animal and vegetal poles. Scores were recorded for each individual in the XIBBIT database. To determine whether egg quality is a stable individual trait in *Xenopus laevis*, we calculated the intraclass correlation coefficient (ICC(1,1)) using a one-way random-effects model. The analysis was restricted to a balanced dataset of 32 frogs, each with exactly three valid quality scores (total N=96 observations).

The ICC was calculated as:

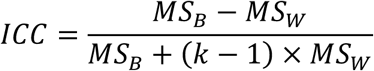

Where:

- MS (Between-frog mean square): Calculated as 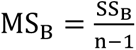, derived from the summed squared deviations of individual frog means from the grand mean.
- MS (Within-frog mean square): Calculated as 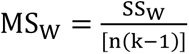, derived from the summed squared deviations of each score from its respective frog mean.
- k: Number of repeated measurements per individual (3).

The significance of the between-frog variance was tested with the F-statistic F = MS_B_/MS_W_, with df_1_ = n − 1 = 31 and df_2_ = n(k − 1) = 64 degrees of freedom.

The 95% confidence interval of the ICC was derived from the F-distribution, and ICC values were interpreted according to the benchmarks proposed by Koo and Li(27): values below 0.50 indicate poor reliability, 0.50-0.75 moderate reliability, 0.75-0.90 good reliability, and above 0.90 excellent reliability.

## Acknowledgements

We would like to express our gratitude to Fabian Prins and Dr. George Azzopardi for developing the original algorithm which is fundamental to this study. We are thankful to all the members of Kamenz lab for their valuable support and feedback using various devices. We also extend our appreciation to the animal caretakers at the animal facility of Faculty of Science and Engineering of the University of Groningen.

---

**Fig S1.**
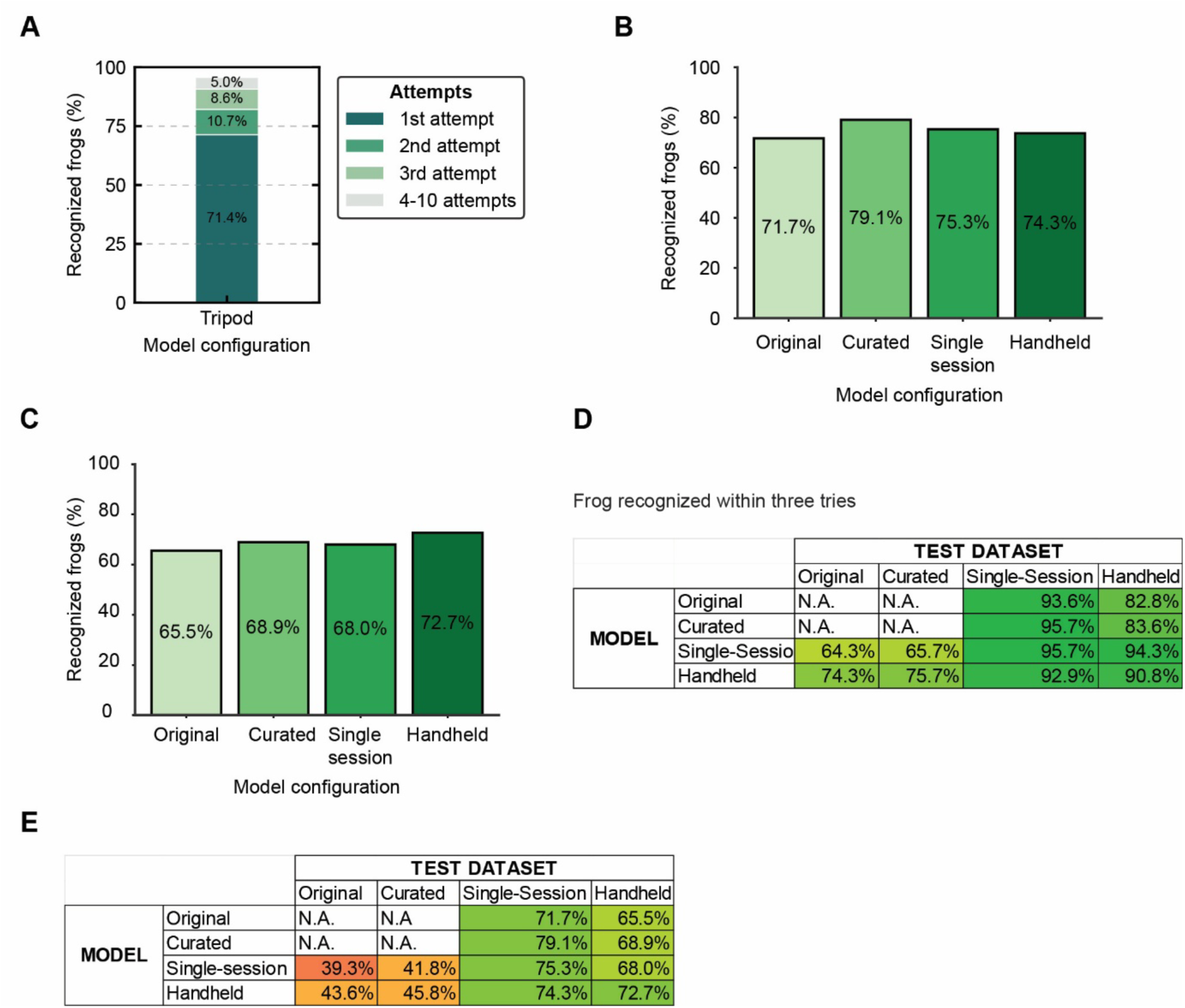
Validation experiment and recognition efficiency summary. **(A)** Cumulative bar chart showing the number of attempts required to correctly identify a frog using the original-trained model tested against an independent tripod-acquired dataset. **(B)** Overall recognition efficiency of all model configurations tested against a single-session tripod-acquired dataset. **(C)** Overall recognition efficiency of all model configurations tested against the single-session handheld dataset. **(D)** Summary table of the percentage of frogs recognized within three attempts across all model configurations (rows) and test datasets (columns); darker shading indicates higher recognition efficiency. **(E)** Summary table of the overall recognition efficiency across all model configurations (rows) and test datasets (columns); darker shading indicates higher recognition efficiency.

**Fig S2.**
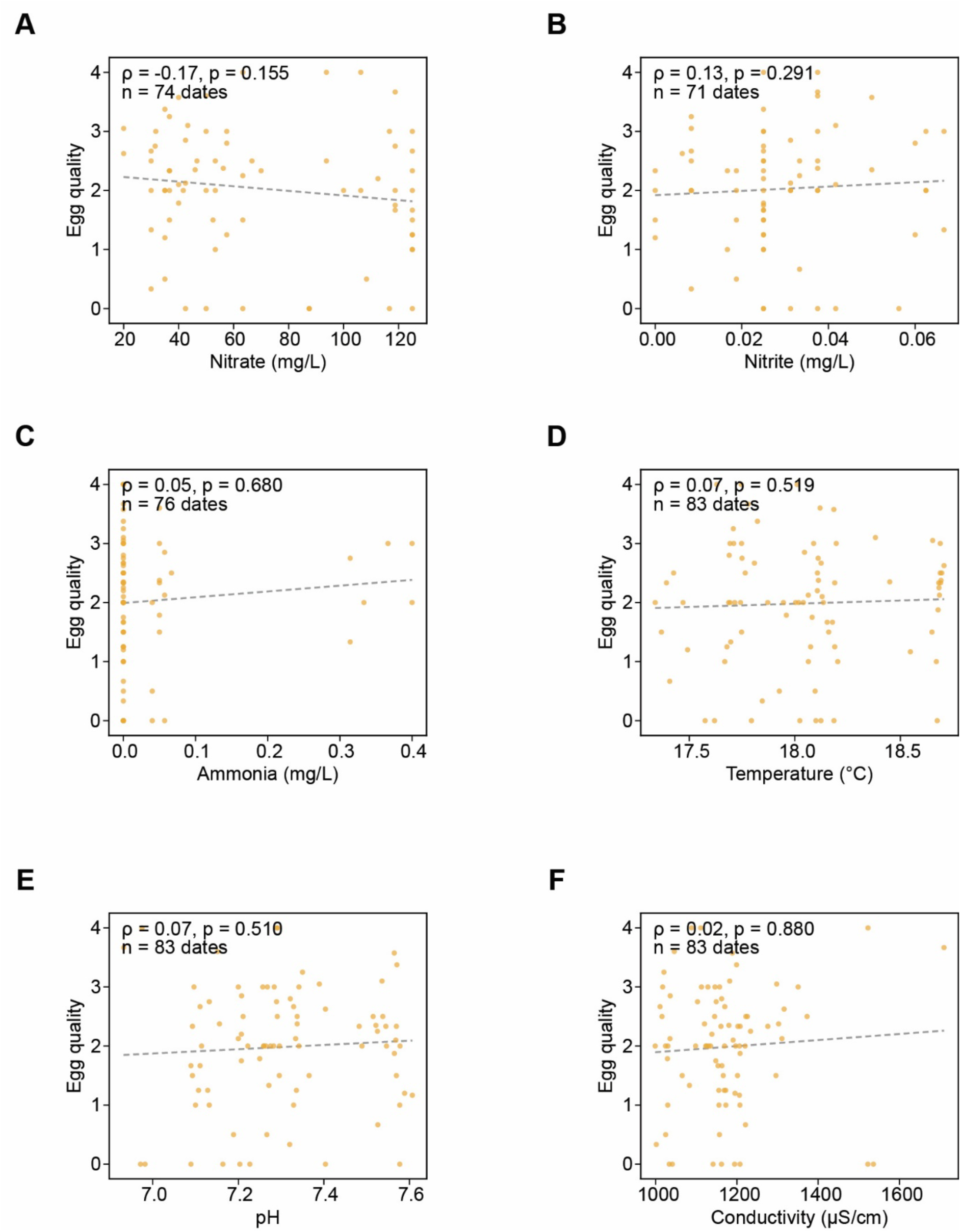
Egg quality is independent of aquatic system water parameters. Scatter plots of egg quality (0–4 scale) against six monitored water-quality parameters: **(A)** nitrate (NO₃, mg/L), **(B)** nitrite (NO₂, mg/L), **(C)** ammonia (NH₄, mg/L), **(D)** temperature (°C), **(E)** pH, and **(F)** conductivity (µS/cm). Each point corresponds to an individual egg-laying date, plotted against the matched water-parameter value; the dashed line indicates the linear trend. For every panel, Spearman’s rank correlation coefficient (ρ), the associated *p*-value, and the number of laying dates with paired measurements (n) are reported in the panel. None of the parameters correlated significantly with egg quality.

